# Spontaneously produced lysogenic phages are an important component of the soybean *Bradyrhizobium* mobilome

**DOI:** 10.1101/2022.05.06.490963

**Authors:** Prasanna Joglekar, Barbra D. Ferrell, Tessa Jarvis, Kona Haramoto, Nicole Place, Jacob T Dums, Shawn W. Polson, K. Eric Wommack, Jeffry J. Fuhrmann

**Affiliations:** Department of Biological Sciences, University of Delaware, Newark, Delaware, USA; Department of Plant and Soil Sciences, University of Delaware, Newark, Delaware, USA; Delaware Biotechnology Institute, University of Delaware, Newark, Delaware, USA; Center for Bioinformatics and Computational Biology, University of Delaware, Newark, Delaware, USA

**Keywords:** Soybean bradyrhizobia, spontaneously induced lysogenic phages, horizontal gene transfer

## Abstract

The ability to nodulate and fix atmospheric nitrogen in soybean root nodules makes soybean *Bradyrhizobium* spp. (SB) critical in supplying humanity’s nutritional needs. The intricacies of SB-plant interactions have been studied extensively; however, bradyrhizobial ecology as influenced by phages has received somewhat less attention even though these interactions may significantly impact soybean yield. In batch culture four SB strains, S06B (*B. japonicum*, S06B-Bj), S10J (*B. japonicum*, S10J-Bj), USDA 122 (*B. diazoefficiens*, USDA 122-Bd), and USDA 76^T^ (*B. elkanii*, USDA 76-Be), spontaneously (without apparent exogenous chemical or physical induction) produced phages throughout the growth cycle; for three strains, phage concentrations exceeded cell numbers by ca. 3-fold after 48 h incubation. Observed spontaneously produced phages (SPP) were tailed. Phage terminase large-subunit protein phylogeny revealed possible differences in phage packaging and replication mechanisms. Bioinformatic analyses predicted multiple prophage regions within each SB genome preventing accurate identification of SPP genomes. A DNA sequencing approach was developed that accurately delineated the boundaries of four SPP genomes within three of the SB chromosomes. Read mapping suggested that the SPP are capable of transduction. In addition to the phages, bacterial strains S06B-Bj and USDA 76-Be were rich in mobile elements consisting of insertion sequences (IS) and large, conjugable, broad host range plasmids. The prevalence of SPP along with IS and plasmids indicate that horizontal gene transfer likely plays an outsized role in SB ecology and may subsequently impact soybean productivity.

**Importance:** Previous studies have shown that IS and plasmids mediate horizontal gene transfer (HGT) of symbiotic nodulation (nod) genes in SB; however, these events require close cell to cell contact which could be limited in soil environments. Bacteriophage assisted gene transduction through spontaneously produced prophages could provide stable means of HGT not limited by the constraints of proximal cell to cell contact. Phage mediated HGT events could be important in SB population ecology with concomitant impacts on soybean agriculture.

## Introduction

The global prominence of soybean, a protein-rich legume, has steadily grown because of its agricultural, economic, and environmental importance in supplying humanity’s protein needs (1). Soybean bradyrhizobia (SB), soybean root-nodulating symbiotic bacteria, transform atmospheric nitrogen (N) into ammonia (NH_3_) (2) providing up to 75% of the plants nitrogen needs (3, 4) and reducing the need for nitrogen fertilization. Promoting biological nitrogen fixation over commercial fertilization also reduces excess nitrogen release to ground and surface waters that can cause eutrophication, as well as, the production of nitrogen gases that contribute to global warming (5). Recent demand for plant-based protein alternatives has further increased the need for efficient and sustainable soybean production (6). Plant-based protein production releases less atmospheric carbon than animal protein production (7, 8) while maintaining similar calorific value (6).

SB strains can differ in their symbiotic effectiveness, *i.e.* their ability to nodulate and subsequently fix nitrogen for the host plant, which in turn depends on a strain’s symbiotic gene repertoire (9, 10). Usually, SB strains having high symbiotic effectiveness are applied to a soybean field to colonize plants with the efficient strain. However, inoculant strains must compete with a diversity of autochthonous SB strains (11). While autochthonous strains may be less effective initially, previous studies have reported the development of soybean-nodulating allochthonous populations (SNAPs) (12) that impact soybean yields. The development of competing SB within soybean fields may correlate with autochthonous soil SB diversity and the proclivity of strains for horizontal gene transfer.

The University of Delaware *Bradyrhizobium* Culture Collection (UDBCC), containing 340 environmental isolates and 12 USDA reference strains of SB having extensive phenotypic and genotypic characterization (13), was established for studying SB ecology and population biology. Interestingly, some UDBCC isolates produce phages spontaneously, *i.e.,* without apparent exogenous chemical or physical induction in batch culture. While several studies have focused on plant-SB interactions, limited data are available on the lysogenic phages of bradyrhizobia (14) and their prophages (15–17).

The evolution of bacteria and phages has been intertwined throughout the history of life on Earth (18). Several biological mechanisms exist within bacteria and phages for neutralizing one another; however, phage-host interactions have also led to the development of mutually beneficial alliances (19). Approximately 10^23^ lytic/lysogenic phage infections occur every second globally (20). As a consequence of these infections, phages can drive microbial evolution by performing genomic rearrangements (21), altering host community dynamics through lysis of dominant populations (22), contributing to bacterial fitness by carrying auxiliary metabolic or virulence genes (23), and participating in horizontal gene transfer (HGT) via specialized or generalized transduction (24). In addition to lytic and lysogenic interactions, several bacteria are known to produce phages spontaneously. *Salmonella* co-culture experiments demonstrated a beneficial alliance where spontaneous induction promoted lysogenic conversion of non-lysogenic strains, ensured propagation of phage DNA, and provided competitive fitness advantage to lysogens by killing sensitive strains (25). The Gram positive bacteria *Lactobacillus gasseri*, a commensal of the human gastrointestinal tract and vagina, has also demonstrated HGT via spontaneously induced phages (26).

In addition to bacteriophages, mobile genetic elements such as plasmids and insertion sequences (IS) play a role in SB biology and ecology. Insertion sequence-mediated HGT and genomic rearrangements have been observed in SB (27, 28). IS sequences work in tandem with large, self-replicating, and conjugable plasmids of rhizobia and bradyrhizobia (29). These mobile elements, in addition to HGT, are involved in recombination, rearrangements, and streamlining of bacterial (30) and SB genomes (31) .

In this study, four UDBCC bacterial strains demonstrating spontaneous prophage production in batch culture were sequenced and assembled. Cultures were monitored for spontaneous phage production, and subsequently the phylogenetic and morphological diversity of these phages was assessed. A sequence read mapping approach was developed enabling unambiguous identification of integrated prophages and the possible involvement of these prophages in HGT. Other genomic analyses focused on identifying genetic elements responsible for HGT in SB. These data were assessed within the context of SB population biology leading to greater appreciation of genomic plasticity within this critically important bacterial genus.

## Results

### Growth and phage production rates vary in studied SB strains

Four UDBCC SB strains demonstrated different growth rates in culture (Fig. 1) based on direct counts with phase contrast microscopy. S06B (*B. japonicum*, S06B-Bj) was the slowest growing strain with a doubling time of 13.2 ± 0.5 h, while the other *B. japonicum* accession, S10J (S10J-Bj), grew nearly twice as fast with a doubling time of 7.1 ± 0.5 h. USDA 122 (*B. diazoefficiens*, USDA 122-Bd) showed a doubling time of 8.9 ± 1.0 h. USDA 76 (*B. elkanii*, USDA 76-Be) was the fastest growing strain with a doubling time of 5.8 ± 1.0 h.

**Figure 1:**
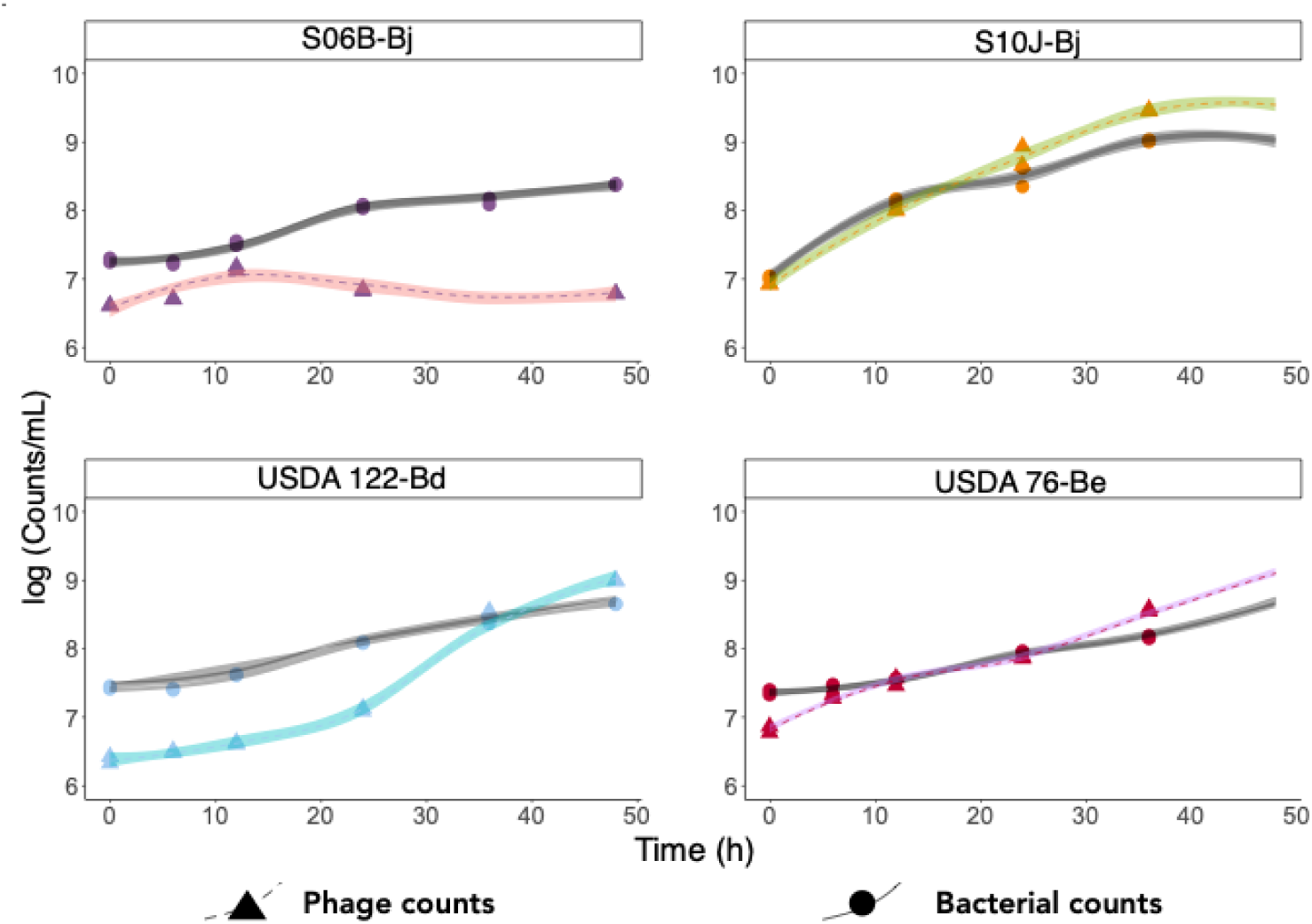
Phage production via spontaneous induction during growth of SB in cell culture. SB cells (filled circles, grey LOESS line) and phage-like particles (filled triangles, colored LOESS line) were counted in duplicate cultures over 48 h. Phage production increased with time for all SB cultures except S06B-Bj.

Phage-like particles (PLPs) were stained with SYBR Gold and enumerated using epifluorescence microscopy. The PLP-to-bacteria ratio increased with time in S10J-Bj, USDA 122-Bd, and USDA 76-Be cultures, and reached a maximum of 3.8, 2.2, and 2.9, respectively, indicating that phage production rate increased with bacterial growth. While PLPs were observed in S06B-Bj culture, the PLP-to-bacteria ratio remained below 0.5 throughout the experiment and decreased to 0.04 at the end of 48 h.

### Tailed phages are spontaneously produced in SB cultures

All spontaneously produced phages observed with TEM were tailed (Fig. 2). Both S06B-Bj and S10J-Bj produced phages with short tails, characteristic of the podovirus-like phage. USDA 122-Bd phages were siphovirus-like with icosahedral capsids and long, noncontractile tails. USDA 76-Be was polylysogenic, producing two different phages, one siphovirus-like phage with long contractile tail and another podovirus-like phage with a short tail. Siphovirus-like phages were more frequent in USDA 76-Be than the podovirus-like phages, however quantitative microscopic enumerations were not performed. All observed phages had head capsid diameters of ∼60 nm (Table 1) suggesting a genome size of approximately 45–60 kb (32).

**Figure 2:**
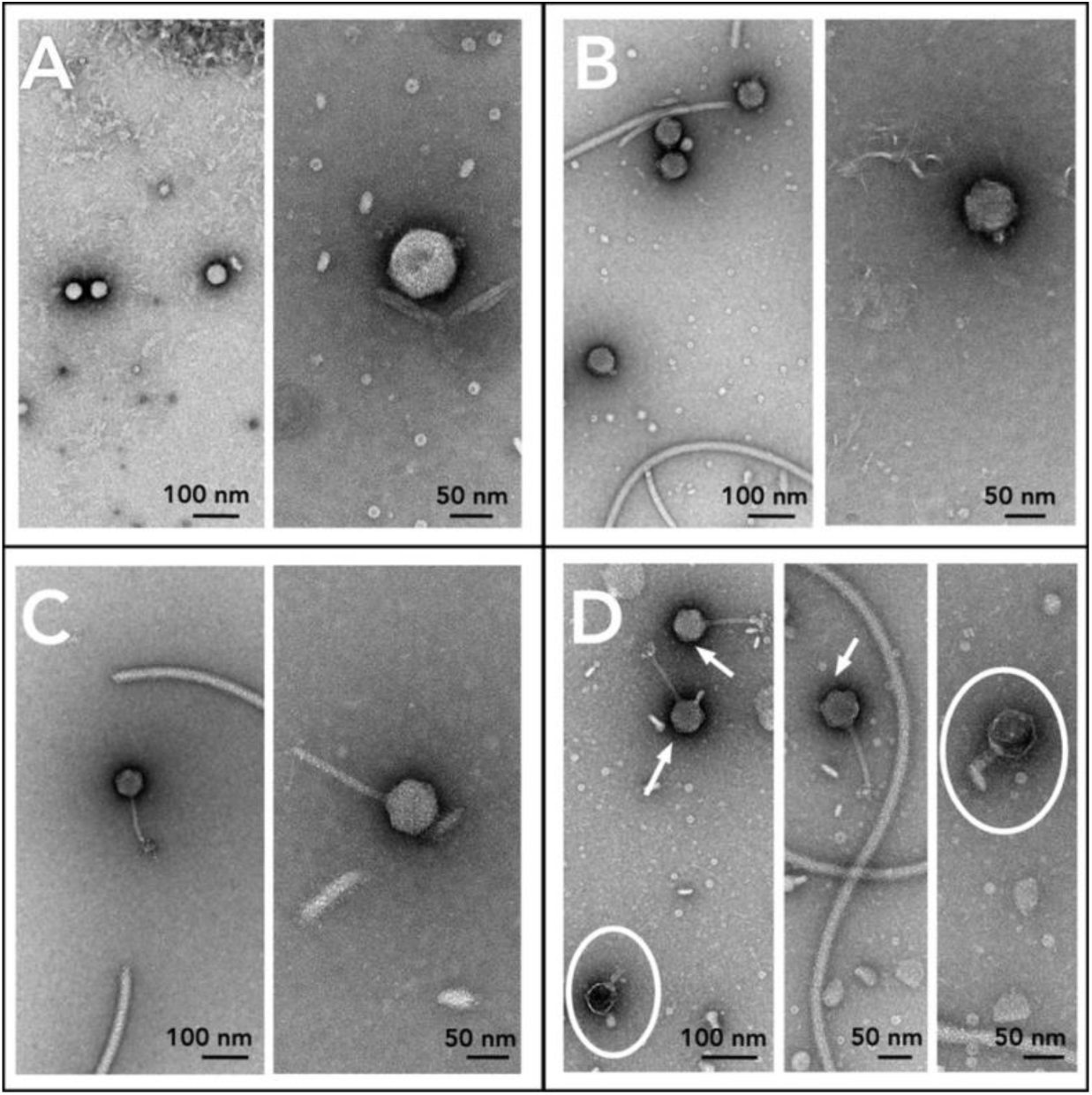
Multiple spontaneously produced phage morphologies observed in SB cultures. Negatively stained (2% uranyl acetate or 1% phosphotungstic acid) transmission electron microscopy images of spontaneously produced phages from SB strains. A) S06B-Bj (podovirus-like), B) S10J-Bj (podovirus-like), C) USDA 122-Bd (podovirus-like), and D) USDA 76-Be (podovirus-like, arrow; and podovirus-like, white circle).

**Table 1:**
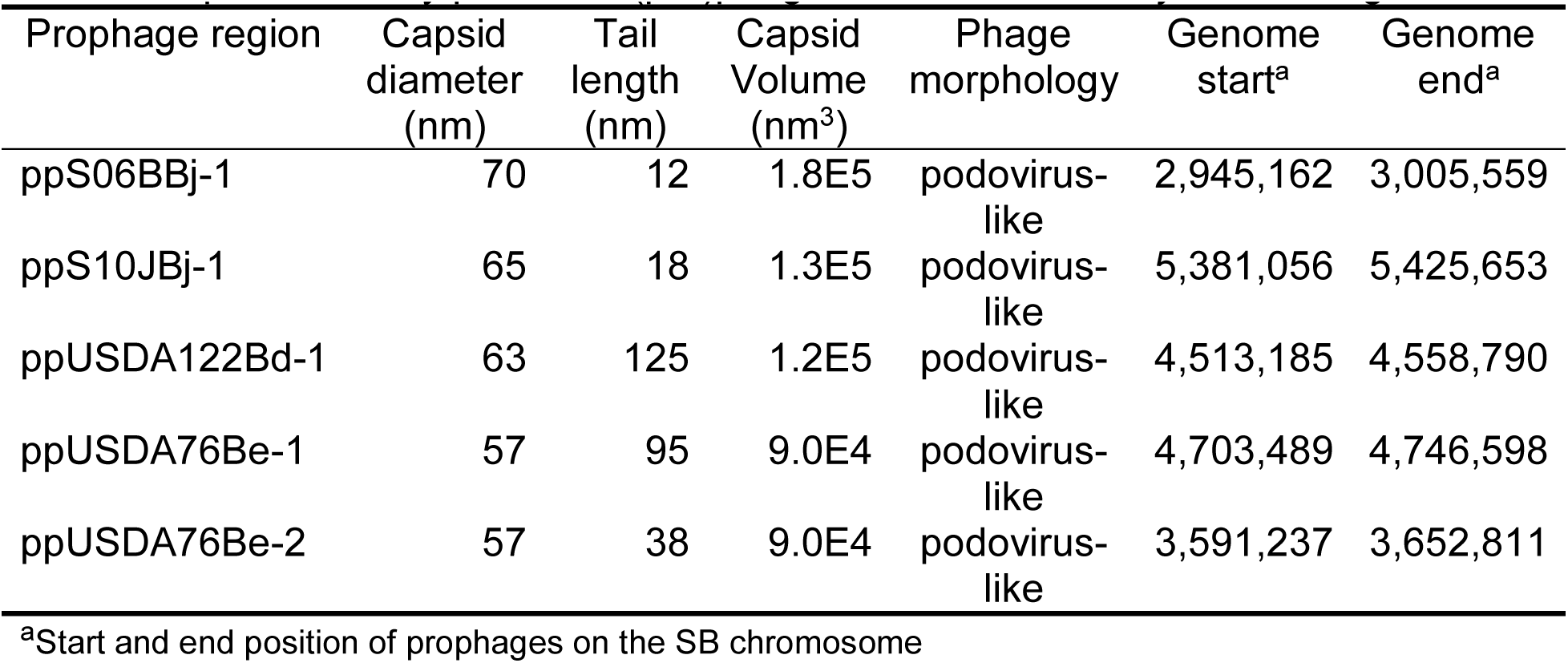
Spontaneously produced (pro)phages identified in *Bradyrhizobium* genome

### Multiple ribosomal operon internal transcribed sequence variants and complete symbiotic islands occur in SB genomes

Bacterial genomes for the studied SB strain ranged between 9-10 Mb (Table 2). In addition to the main chromosomal contig, S06B-Bj and USDA 76-Be genomes contained three and one plasmid, respectively (Fig. 3, Fig. 4, Table 2). Analysis of conserved marker genes by CheckM analysis indicated the genomes were 100% complete with possible contamination levels of 0.18– 0.34%, well within acceptable levels to be considered complete genomes. Previous PCR sequencing studies showed the presence of three different ITS variants in S10J-Bj genome (13). Although the final S10J-Bj genome assembly only contained two copies of single ITS variant, mapping of error-corrected PacBio S10J-Bj reads (Supplementary Fig. S1) supported the existence of three heterogeneous ITS variants (13).

**Figure 3:**
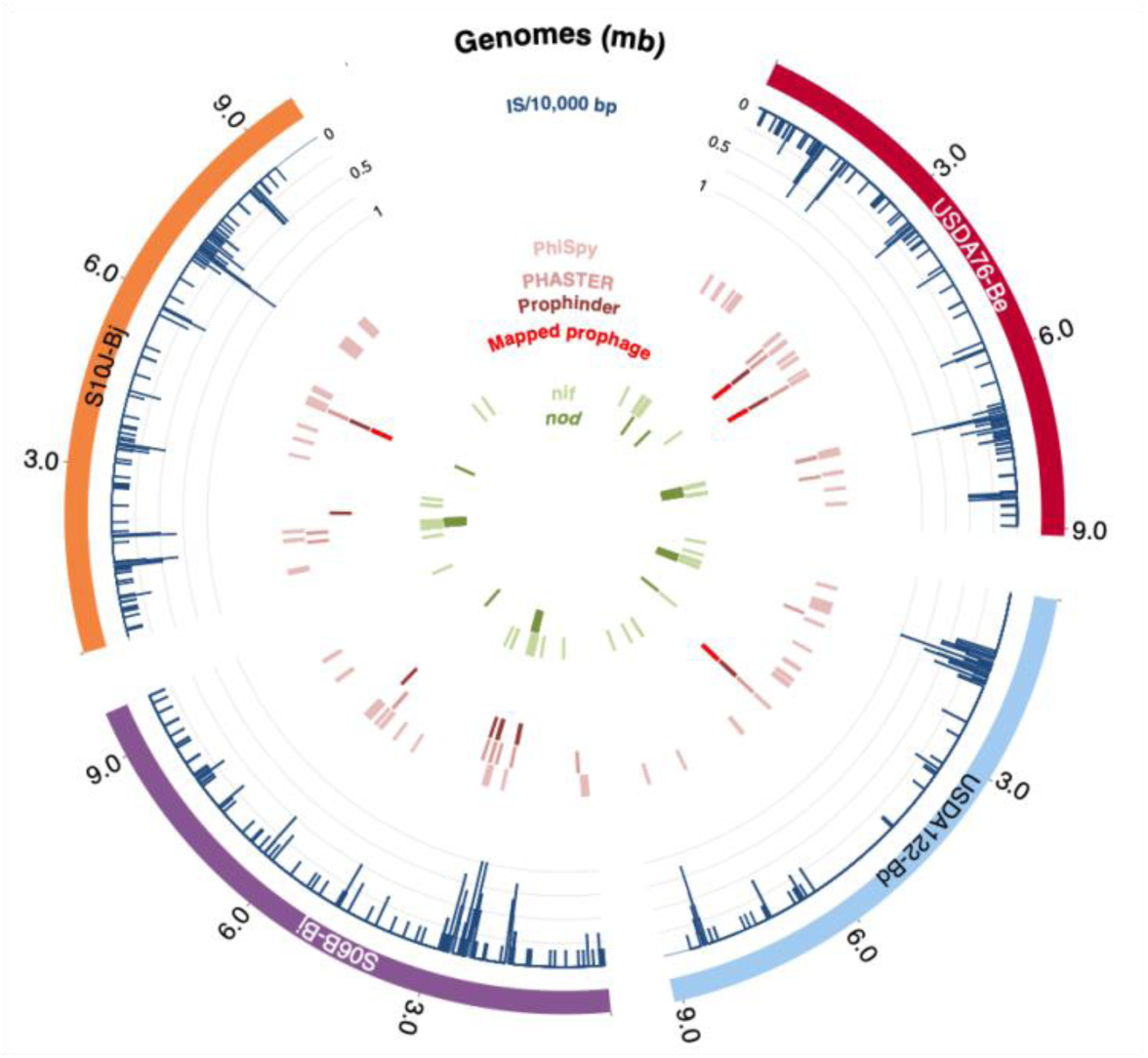
Chromosomal annotations of *Bradyrhizobium* genomes. Each genome is represented on the outermost ring. The density of insertion sequences per 10,000 bp is represented by blue lines as a proportion of the maximum density. Additional rings show the chromosomal locations of prophage predictions from PhiSpy, PHASTER, and Prophinder, as well as the sequence-mapped prophage regions. The two innermost rings show the location of nitrogen fixation (*nif*) and nodulation (*nod*) genes.

**Figure 4.**
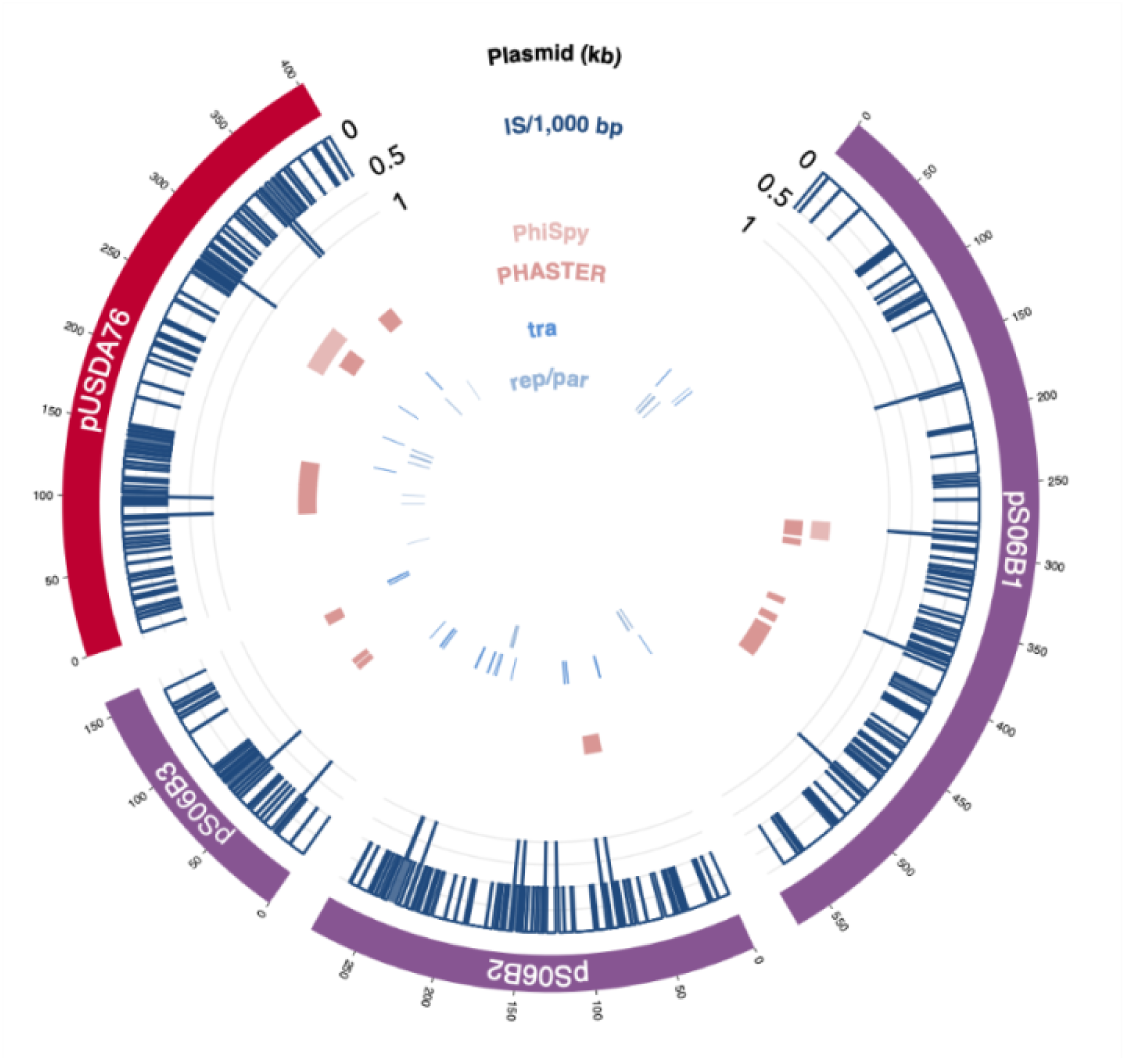
Annotations of *Bradyrhizobium* plasmids. Plasmids from S06B-Bj and USDA 76-Be are represented on the outermost ring. The density of insertion sequences per 1,000 bp is represented by blue lines as a proportion of the maximum density, followed by rings showing the genomic locations of prophage predictions from PhiSpy and PHASTER. The two innermost rings show the location of conjugation genes (*tra*) and replication/partition genes (*rep*/*par*) genes.

**Table 2:** *Bradyrhizobium* genome assembly metrics

| UDBCC<sup>a</sup><br>accession | Species | Genetic<br>element | NCBI<br>accession | Length<br>(bp) | Coverage<br>(X) | Reads<br>(#) | GC<br>(%) | tRNA<sup>b</sup><br>(#) | rRNA<sup>c</sup><br>(#) | CDS<sup>d</sup><br>(#) |
| --- | --- | --- | --- | --- | --- | --- | --- | --- | --- | --- |
| S06B-Bj | *B. japonicum* | Chromosome | CP066351 | 9,745,549 | 90 | 109,402 | 63.5 | 56 | 6 | 9521 |
|  |  | pS06B1 | CP066352 | 572,704 | 121 |  | 60.1 | 4 | 0 | 638 |
|  |  | pS06B2 | CP066353 | 279,984 | 130 |  | 61.2 | 0 | 0 | 450 |
|  |  | pS06B3 | CP066354 | 159,206 | 100 |  | 61.7 | 0 | 0 | 183 |
| S10J-Bj | *B. japonicum* | Chromosome | CP181350 | 9,864,868 | 250 | 214,517 | 63.5 | 56 | 6 | 9441 |
| USDA 122-Bd | *B. diazoefficiens* | Chromosome | CP066355 | 9,137,248 | 100 | 70,204 | 64.0 | 50 | 3 | 8619 |
| USDA 76-Bd | *B. elkanii* | Chromosome | CP066356 | 9,115,474 | 210 | 216,943 | 63.9 | 47 | 3 | 8661 |
|  |  | pUSDA76 | CP066357 | 402,731 | 150 |  | 60.4 | 0 | 0 | 485 |

<sup>a</sup> University of Delaware *Bradyrhizobium* Culture Collection
<sup>b</sup> tRNA genes
<sup>c</sup> rRNA operons

Each genome contained a complete set of nodulation and nitrogen fixation genes. USDA 76-Be contained a ∼9 kb complete rhizobitoxine (*rtx*) island consisting of *rtxACDEFG* genes (Supplementary Fig. S2), consistent with other *B. elkanii* strains that produce rhizobitoxine, an enol ether amino acid that initially promotes nodulation but causes foliar chlorosis in sensitive soybean cultivars (33, 34). Interestingly, S06B-Bj and S10J-Bj, both *B. japonicum* species not known for producing rhizobitoxine, carried a contiguous ∼9 kb region with individual ORFs 70–80% identical (nucleotide) to the *rtx* operon (Supplementary Fig. S2). USDA 122-Bd did not encode any *rtx* genes.

### SB mobilome is diverse and varied across species

SB with large numbers of IS (insertion sequences) are called highly reiterated sequence (HRS) strains (31). HRS strains have reduced growth rates and an increased capacity for horizontal gene transfer (HGT) (35). The distribution of IS was used for assessing the potential for gene rearrangement and HGT in each SB genome.

ISEscan predicted 151 and 152 IS in S10J-Bj and USDA 122-Be (Table 3), respectively, representing ca. 2% of the total genome. A total of 410 and 276 IS, accounting for ∼4–6% of the genome, were predicted in S06B-Bj and USDA 76-Be respectively, a characteristic of HRS strains (31). IS density was higher near nodulation and nitrogen fixation genes (Fig. 3). Three major classes of IS were identified in SB (Fig. 5): 1) DDE type (active site contains two aspartic acid (D) and one glutamic acid (E)), 2) DEDD type (single IS family, IS110; active site contains one aspartic acid, one glutamic acid, and two additional aspartic acids), and 3) HUH type (single IS family, IS91; histidine-large hydrophobic amino acid-histidine) (36). IS from family ISCNY have not yet been classified into a particular type. The DDE type, which contains 16 different IS families, was the most abundant. IS110 (single DEDD type family) was one of the five most abundant IS families. While a previous study had reported 1–6 copies of IS110 (31) in HRS strains, the SB sequenced in this study carried 2–25 copies of this IS. Consecutive duplicates of IS21, IS66, IS630, and IS110 elements were integrated 93, 30, 28, and 66 times, respectively.

**Figure 5.**
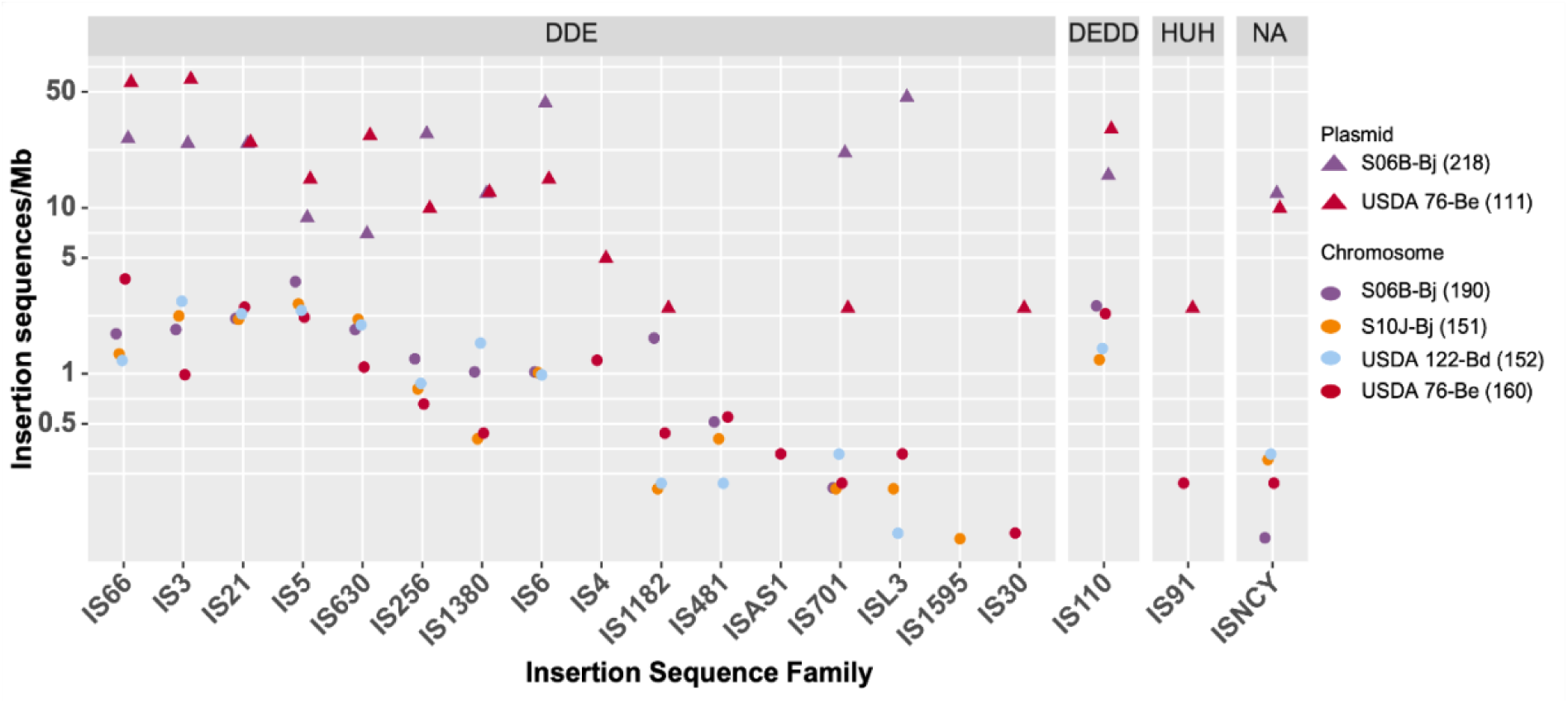
Plasmids contribute a large proportion of insertion sequences in SB strains. Log normalized abundance of IS families per 1 Mb is provided for each SB chromosome (filled circle) and the plasmids (filled triangle). The total number of IS for each chromosome or collection of plasmids are reported in parentheses. IS are grouped by the three major classes of IS families: DDE (active site containing two aspartic acid (D) and one glutamic acid (E)); DEDD (active site containing one aspartic acid, one glutamic acid, and two additional aspartic acids), 3) HUH type (histidine (H) - large hydrophobic amino acid (U) - histidine), and 4) and 4) NA (not yet been classified into a particular type; includes IS from ISCNY family).

**Table 3:** Insertion sequences in *Bradyrhizobium* genomes and type strains

| Organism | Species | Insertion sequences (#) | Insertion sequence density (#/Mb) |
| --- | --- | --- | --- |
| S06B-Bj | *B. japonicum* | 410 | 40 |
| S10J-Bj | *B. japonicum* | 151 | 15 |
| NK6 | *B. japonicum* | 560 | 55 |
| USDA 6<sup>b</sup> | *B. japonicum*<sup>a</sup> | 120 | 13 |
| USDA 122-Bd | *B. diazoefficiens* | 152 | 16 |
| USDA 110<sup>b</sup> | *B. diazoefficiens*<sup>a</sup> | 104 | 12 |
| USDA 76-Be | *B. elkanii* | 276 | 31 |
| K12-MG1655<sup>b</sup> | *E. coli*<sup>a</sup> | 60 | 13 |
| <sup>a</sup>Type strains |  |  |  |
| <sup>b</sup>IS previously identified in NCBI accession |  |  |  |

Several IS identified in the S06B-Bj and USDA 76-Be genomes were present on plasmids (Fig. 4). Comparison against PLSDBs plasmid database (37) suggested these large plasmids were similar to those found in other *Bradyrhizobium* spp. (Table 4). A *repABC* operon, partition, and conjugation genes were identified on each of the plasmids (Fig. 4). Additionally, the plasmids contained metabolic genes including serine dehydratase, serine hydroxymethyl transferase, class III ribonucleotide reductase, and some nodulation proteins (data not shown).

**Table 4:** PLSDB plasmid similarity to sequenced *Bradyrhizobium* plasmids

| NCBI accession | Species | Average identity <sup>a</sup> |  |  |  |
| --- | --- | --- | --- | --- | --- |
|  |  | pS06B1<br>(*B. japonicum*) | pS06B2<br>(*B. japonicum*) | pS06B3<br>(*B. japonicum*) | pUSDA76<br>(*B. elkanii*) |
| NZ_CP049700.1 | Unclassified<br>sp. 323S2 | 0.95 (3.15) | - | - | 0.96 (441) |
| NZ_AP014686.1 | *B. diazoefficiens* | 0.93 (244) | 0.95 (330) | 0.90 (110) | 0.92 (178) |
| NZ_AP014687.1 | *B. diazoefficiens* | 0.90 (110) | - | 0.95 (315) | - |
| NZ_AP014659.1 | *B. diazoefficiens* | - | - | 0.93 (244) | - |

<sup>a</sup>Data generated from PLSDB, p value: 0.1, percent identity cutoff 0.9, winner takes all strategy

### Inconsistencies between prophage prediction algorithms

PHASTER, Prophinder, and PhiSpy identified putative prophages on all SB chromosomes and many of the plasmids. However, the number and size of prophage regions predicted by these algorithms varied greatly.

PHASTER identified a total of 31 potential prophages (12 incomplete, 8 intact, 11 questionable), 14 of which were found on plasmids (Fig. 3, Fig. 4). Overall, the predicted prophage sizes (7–35 kb) (Table 5) were smaller than those estimated from phage capsid volumes (45–60 kb). BLASTp analysis revealed that ∼80% of the PHASTER-predicted prophage regions contained IS21 and IS5 families and hypothetical proteins. Prophinder predicted nine prophage regions ranging 10-45 kb in size, all on chromosomes. PhiSpy predicted a total of 65 potential prophages with genome sizes of 2–106 kb with two prophages predicted on plasmids. Like PHASTER, many of the PhiSpy predicted prophages contained insertion sequences and hypothetical proteins. Only a few predicted prophage regions were common across the tools (e.g, Fig. 6). Even in these cases, the predictions were small with inconsistent boundaries. Thus, an alternative sequencing and mapping approach was designed for delineating the prophage regions.

**Figure 6:**
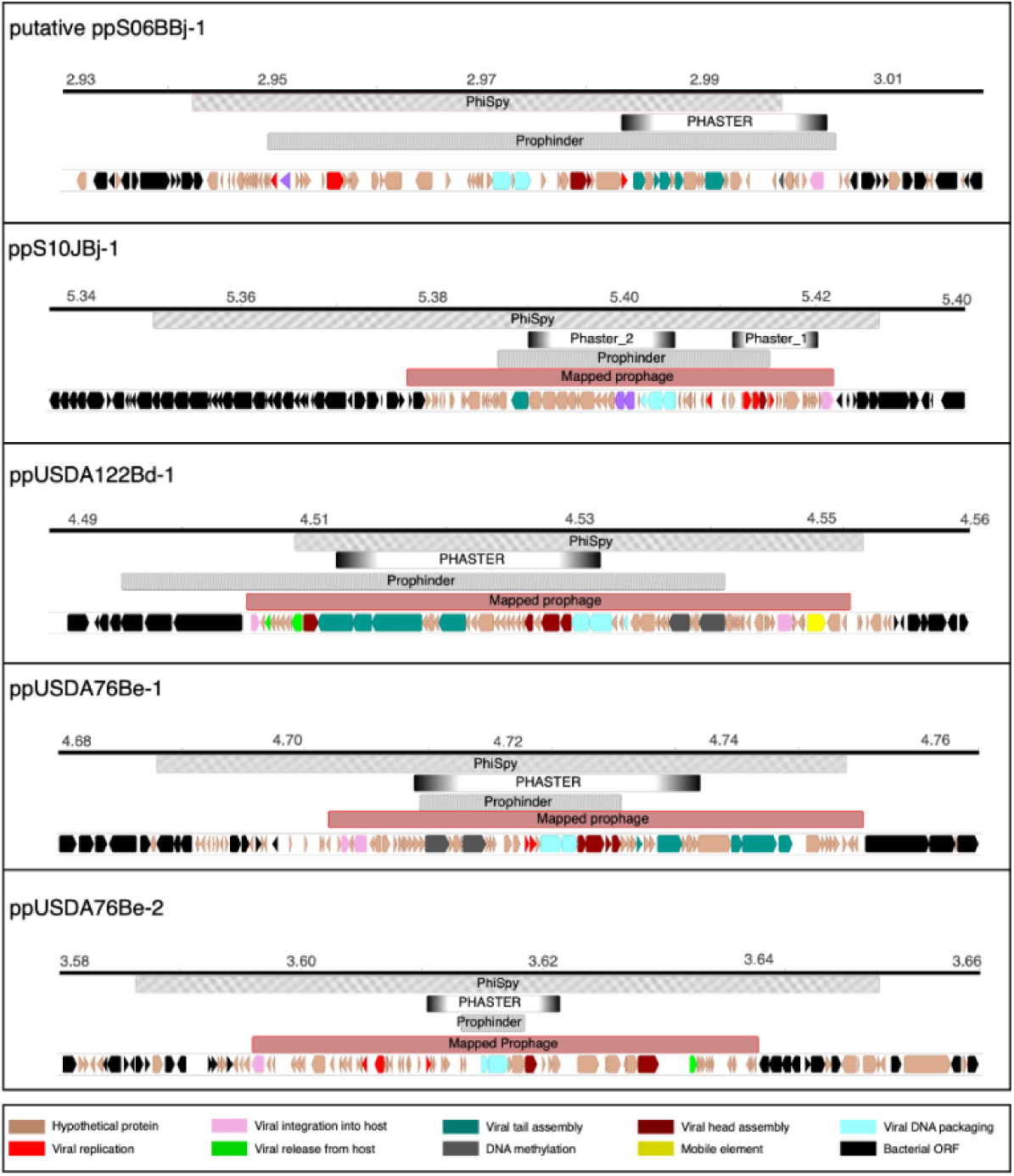
Phage boundaries predicted by bioinformatic tools and mapping of sequencing reads. Bioinformatic software PhiSpy, PHASTER, and Prophinder predict inconsistent and often incomplete prophage regions. Mapping of sequenced spontaneously produced phage (red) delineates prophage boundaries congruent with phage genome size predictions based on TEM images of phage observed in SB cultures. The predicted ORFs from mapped prophage regions were BLASTed against UniRef and the NCBI Virus database to assign functional annotations. The ORFs are colored according to their functional GO Terms.

**Table 5:** Prophage regions predicted by bioinformatic tools

| Genome | PHASTER | PhiSpy | Prophinder |
| --- | --- | --- | --- |
| S06B-Bj | 17 | 14 | 3 |
| S10J-Bj | 4 | 18 | 2 |
| USDA 122-Bd | 3 | 18 | 1 |
| USDA 76-Be | 7 | 14 | 2 |
| Prediction sizes (kb) | 11–35 | 2–106 | 10–45 |

### Phage read mapping accurately delineated prophage boundaries

Sequencing of phage DNA isolated from SB cultures yielded between 8 and 10 million reads for the viral fraction in each experiment. Around 80–94% of the phage reads mapped to the bacterial genome. Read coverage across most of the S10J-Bj, USDA 76-Be, and USDA 122-Bd genomes was ∼35x. However, there were 40–50 kb regions which showed high coverages of between 60,000 and 150,000x (Fig. 7) indicating that these were regions corresponding to spontaneously produced phages (Fig. 2, Table 4). One enriched mapping site was observed in each of the USDA 122-Bd (ppUSDA122Bd-1) and S10J-Bj (ppS10JBj-1) chromosomes, and two sites were observed on the USDA76-Be (ppUSDA 76Be-1 and ppUSDA76Be-2) chromosome, corroborating TEM observations of two different phage morphologies (Fig. 2). Prophage ppUSDA76Be-1 had more coverage (Fig. 7) than ppUSDA76Be-2, indicating the earlier phage was more frequently produced. Furthermore, phage capsid protein phylogeny (data not shown) showed that ppUSDA76Be-1 was a siphovirus-like phage and ppUSDA76Be-2 was a podovirus-like phage. All prophages identified through sequence mapping corresponded with some of the bioinformatic predictions (Fig. 6). However, in no case did the bioinformatic predictions exactly match the read mapping results.

**Figure 7:**
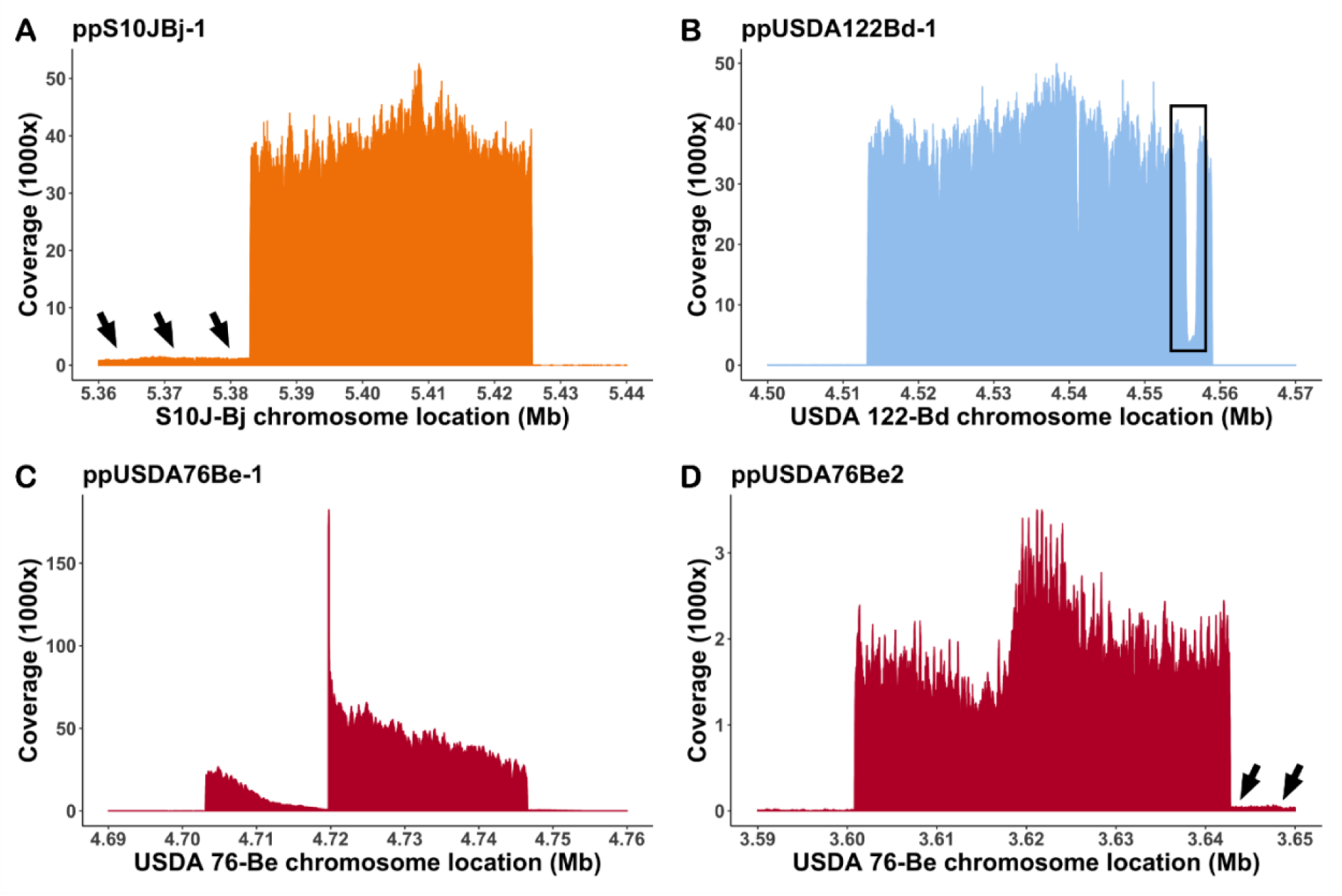
Phage DNA sequence reads mapped to their respective bacterial chromosome showed specific regions of enriched coverage. A single enriched region was observed for A) S10J-Bj and B) USDA 122-Bd, consistent with a single spontaneously produced phage observed within these cultures. The boxed region in panel B indicates an area of low coverage containing an insertion sequence in the USDA122 prophage. Two enriched regions were observed in USDA 76-Be (C & D), consistent with the two phage morphologies observed in culture. Adjacent regions of moderately increased coverage (black arrows) were observed A) upstream of ppS10JBj-1 and D) downstream of the ppUSDA76Be-2 suggesting that these genome regions may also be packaged in phage heads.

In addition to the prophage, six other regions of enriched mapping were observed on the USDA 122-Bd chromosome (Supplementary Fig. S3). Examination of gene content underlying these peaks indicated similarity to an insertion sequence present in the USDA 122-Bd prophage. Additionally, regions immediately flanking prophages ppUSDA76Be-2 and ppS10JBj-1 (Fig. 7) containing only bacterial ORFs also showed higher coverage (∼500–2000x) than the rest of the bacterial genome (∼35x), indicating that these flanking sites may have been packaged into phage capsids at a lower frequency than the rest of the prophage. S06B-Bj was a low spontaneous phage producer which resulted in low phage DNA yields (Fig. 1). Mapping of S06B-Bj phage reads did not show enriched mapping and resulted in even coverage across the genome, except for regions containing IS66 insertion sequences (Supplementary Fig. S3). However, one putative prophage site identified by PhiSpy, Prophinder, and PHASTER on the S06B-Bj chromosome also contained a high number of phage related proteins (Fig. 6). Sequence reads from bacterial DNA was also mapped to the bacterial genomes as controls for the phage DNA sequence read mapping experiments. In each case bacterial sequence reads showed even mapping (200–1000x) across the genome (Supplementary Fig. S4).

### Spontaneously induced bradyphages are phylogenetically diverse

The large subunit of the terminase protein, TerL, is responsible for phage genome packaging and identification of the phage genome termini. Three major types of phage genome termination mechanisms, 3’ cohesive ends, 5’ cohesive ends, and headful packaging, can be identified from TerL phylogeny (Fig. 8) (38). Prophage TerL proteins from ppUSDA76Be-1 and ppUSDA122Bd-1 (both siphovirus-like phages) were similar and predicted to have cos 3 packaging, as these sequences clustered with other 3’ cohesive end TerL proteins containing a terminase-1 conserved protein family (Pfam) domain. Prophage TerL proteins from ppS10JBj-1 and ppUSDA76Be-2 prophages (both podovirus-like phages) were predicted to have headful packaging, as these sequences clustered with headful-packaging TerL proteins containing either terminase- 3 or terminase-6 Pfam domains. Lastly, the bioinformatically predicted ppS06BBj-1 prophage (podovirus-like) was predicted to have cos 5 packaging, as its TerL protein clustered with 5’ cohesive-end terminases having a terminase-GpA Pfam domain.

**Figure 8:**
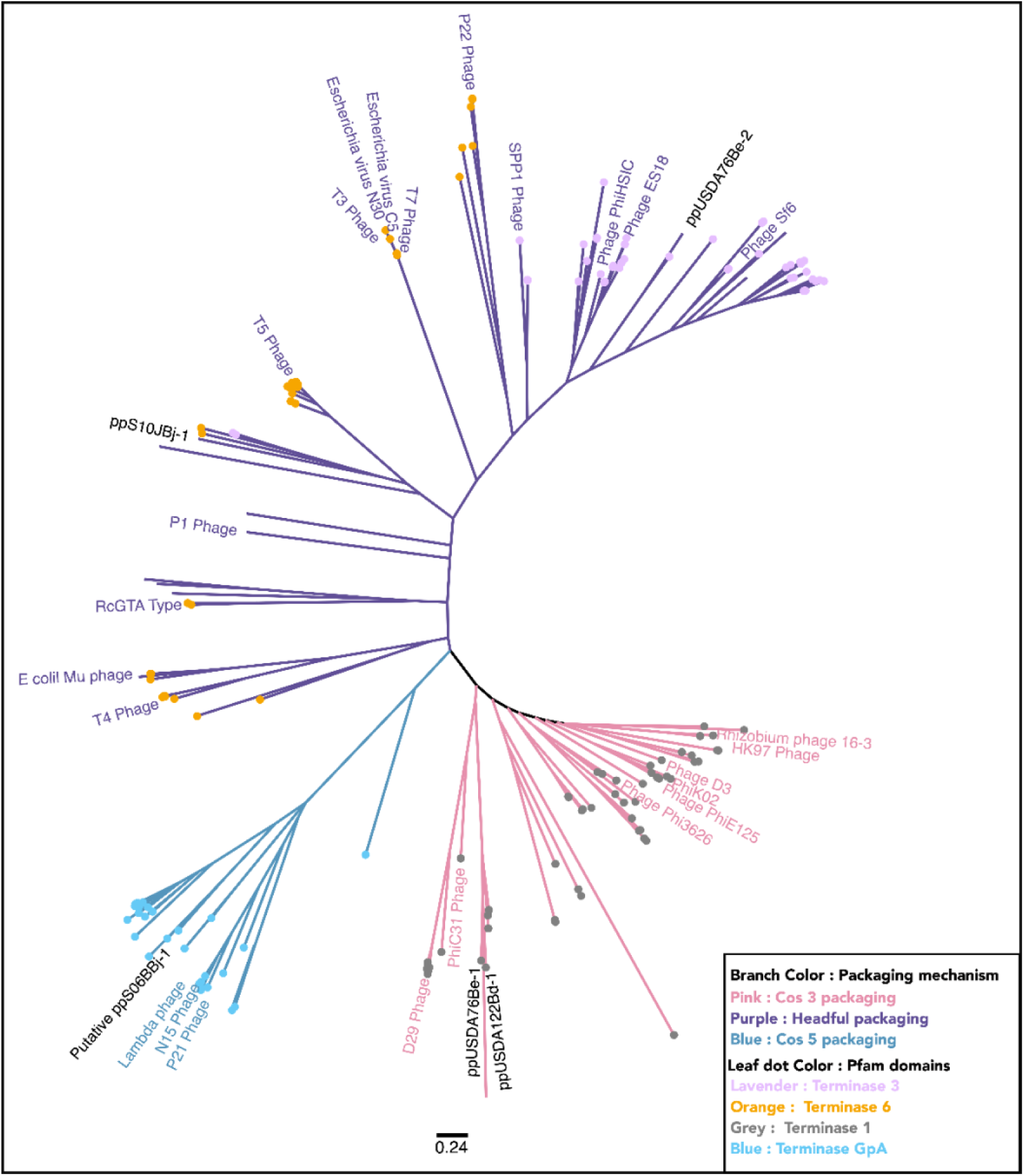
Phylogenetic analysis of large terminase subunit (TerL) proteins indicates that SBl spontaneously produced phages demonstrate a variety of genome packaging and termination types. Unrooted phylogenetic tree of TerL amino acid sequences from SB phages and 242 reference sequences extracted from the UniProt and RefSeq databases shows the diversity of phage packaging mechanisms observed in spontaneously produced bradyphages. Branches are colored according to clades specific for phage genome packaging mechanisms cos 3 type (pink), cos 5 type (blue), and headful packaging (purple) mechanisms based on clade membership of reference sequences with confirmed packaging mechanisms (colored labeled branches). Labels for SB spontaneously produced prophages are shown in black text. Unlabeled branches indicate reference phage TerL sequences for which genome termination type has not been experimentally verified. Leaf tip dots indicate the conserved domains (protein family, Pfam) found in TerL proteins. Scale bar indicates amino acid changes per residue.

## Discussion

### Conflicting evidence for multiple rRNA operons in S10J-Bj genome

Limitations of different sequencing technologies often causes incomplete genome assembly; however genomic rearrangements and tandem repeats can also contribute to incomplete assembly (39). While the assemblies of S06B-Bj, USDA 122- Bd, and USDA 76-Be yielded complete contiguous genomes, the S10J-Bj genome assembly had several complications.

Previous S10J-Bj ITS amplicon sequence analysis showed the presence of three heterogenous ITS sequences (variants 1, 2, and 3) in S10J-Bj genomic DNA (13). While the corrected PacBio S10J-Bj reads showed the presence of all three ITS sequence variants (Supplementary Fig. S1), draft assembles produced by HGAP2 and MaSurCA hybrid assembly (40) contained only ITS sequence variants 2 and 3 in two *rrn* operons. Interestingly, the HGAP4 assembly had two identical copies of ITS variant 3 in both of its *rrn* operons. While the presence of two *rrn* operons in a *B. japonicum* strain is not surprising, the presence of three ITS variants in S10J-Bj with two *rrn* operons is inexplicable. Previous studies have shown that repeated recombination events between *rrn* operons of *Vibrio cholerae* led to the formation of new ITS variants (41), and perhaps such recombination events in S10J-Bj may produce subpopulations with different ITS variants. It is possible that S10J-Bj cultures may be comprised of two or three populations, with ITS variant 3 being the predominant population. Another scenario is that the S10J-Bj culture may be contaminated, but this seems unlikely as DNA for each of the sequencing experiments (ITS amplicon, PacBio genome sequencing, and Illumina sequencing) was taken from a single colony at different timepoints. Additional higher sequencing depth may help resolve conflicting observations regarding these ITS variants.

### Mapping of phage reads resolves inconsistencies of bioinformatic prophage prediction

Several bioinformatic algorithms have been developed for the identification and prediction of prophages in the bacterial genomes. Prophage predictions using three bioinformatic algorithms (PHASTER, Prophinder, and PhiSpy) were inconsistent and incomplete. Phage genomes are modular and typically organized into early, mid, and late phase genes (42), and repeated recombination and mutation events between the different modules (43) can confound bioinformatic predictions. Each algorithm relies on homology to known phage proteins as a key heuristic in predicting prophage regions. However, phages often contain a high proportion of hypothetical proteins and proteins of unknown function (44) creating problems for accurate prophage prediction. PHASTER predicted small prophages that consisted of structural proteins, integrases, and terminases, and a few of these predicted regions corresponded with prophages identified by sequence mapping (Fig. 6). However, PHASTER predictions did not include all hypothetical proteins, probably because these peptides lacked homology to the database utilized by the algorithm. While Prophinder’s predictions were generally larger than PHASTER’s and localized with mapped prophages, the underlying Prophinder database was last updated in 2010 (45) which may have limited accuracy. PhiSpy predicted numerous prophages and overpredicted the length of the mapped prophages. Like PHASTER, PhiSpy too identified many false positives containing only IS and hypothetical proteins. Overall, these results clearly indicate that the three bioinformatic algorithms alone cannot accurately identify or delineate prophages. Understanding of the role of inducible prophages in bacterial population biology requires accurate data. The sequence mapping approach developed in this study was an important step towards improving data accuracy.

Because bioinformatic prophage predictions were highly inaccurate, an alternative experimental approach was developed for prophage identification. In this approach DNA from spontaneously produced phage was isolated after separating phage particles from cells and dissolved DNA. Phage DNA was sequenced alongside bacterial genomic DNA isolated from cultures. Phage DNA sequences were assembled resulting in prophage genomic contigs (data not shown). Phage sequence reads were mapped to the host genomes (Figs. 7, S3) in an approach similar to other recently reported work (46). The assembly and mapping approaches agreed in terms of size, gene orientation, and coverage; however, the mapping analysis provided additional information relevant for phage biology. Mapping of phage reads from USDA 76-Be and USDA 122-Bj showed high coverage regions adjacent to ppUSDA76Be-2 and ppS10JBj-1 (Fig. 7). Coverage differences between the prophage and flanking regions (∼65,000x versus ∼2000x, respectively) likely caused the assembly approach to miss these flanking regions. Given that contaminating DNA was reduced by DNase digestion of the phage concentrates prior to sequencing, the high coverage of the adjacent regions (∼2000X) compared to background DNA coverage (∼35X), likely indicates heterogeneity in phage packaging mechanisms and involvement of these phages in transduction (46). Additionally, a region of decreased coverage was observed in ppUSDA76Be-1 (Fig. 7). Although the reason for this dip is unknown, other authors have suggested that such mapping profiles are indicative of phage packaging mechanisms (47).

Phage reads mapped to the S06B-Bj genome showed even coverage across the genome, except for regions that encoded for insertion sequences (Supplementary Fig. S3). This is reminiscent of small phage-like particles known as gene transfer agents (GTA) that exclusively package bacterial DNA in their capsids (48–50). While the S06B-Bj genome did not contain any GTA-like genes and the phage particles produced in batch culture were larger than previously reported GTA particles (49), it is nevertheless possible that S06B-Bj phages are demonstrating similar behavior to better known GTAs and are packaging bacterial DNA because of rigorous DNase treatment used to reduce bacterial DNA contamination.

### The potential impact of IS elements, plasmids, phages, and horizontal gene transfer on SB population biology

Insertion sequences (IS), plasmids, and bacteriophages constitute the mobilome of bacteria and are often responsible for horizontal gene transfer (HGT) (51). Many IS elements carry genes encoding transposases and related regulatory proteins flanked by inverted repeats which allow for movement of IS within the genome using a copy-paste or cut-paste mechanism (30, 52). Highly repetitive sequence (HRS) SB strains have 250–800 IS elements (31), typically with high copy numbers (150–250 copies/genome) of IS630 (ISRj1), IS3 (ISRj2), and IS1630. The total number of IS elements in the S06B-Bj and USDA 76-Be genomes (410 and 276, respectively) indicated they were HRS strains. However, these strains were not similar to other HRS strains in terms of the copy numbers of specific IS. All observed IS families were evenly distributed within the S06B-Bj and USDA 76-Be genomes (Fig. 5), and the number of IS elements from each IS family ranged from 1-40 per genome. The high number of IS elements in the S06B-Bj and USDA 76-Be genomes also suggested elevated levels of gene transfer and rearrangement activities, indeed previous studies have shown that HRS strains transfer *nod* genes from *B. elkanii* to *B. japonicum* via HGT (28).

The potential impacts of IS elements on SB diversity vary. Many IS can function as composite transposons, formed when two independent IS sequences mobilize the intervening host DNA (52). In addition, many IS observed within the SB genomes were arranged adjacently. In many cases such arrangements provide an active ∼35 bp promoter region increasing transposition activity (30, 52), and leading to greater chances for gene interruption or transposition.

An important IS gene interruption was observed in the rhizobitoxine (*rtx*) operon (*rtxACDEFG* genes) of S06B-Bj (Supplementary Fig. S2). While *rtx*-like genes were reported in *B. diazoefficiens* type strain USDA 110^T^ this strain does not produce rhizobitoxine (53). The *rtxA* gene, which catalyzes the production of dihydrorhizobitoxine necessary for rhizobitoxine production, was truncated by a premature stop codon in USDA 110 ^T^. S06B-Bj, S10J-Bj, and type strain USDA 6^T^ (*B. japonicum*) (54) also carried *rtx*-like regions (Supplementary Fig. S2) with similarly truncated *rtxA* genes. Interestingly, the *rtxA*-like gene in S06B-Bj was truncated by two IS elements. It is possible that the repeated integration of IS played a role in truncation and evolution of the *rtx* operon.

Plasmids are another major part of the SB mobilome that can contribute to HGT (51). The megaplasmids (plasmids ≥ 100 kb (55)) identified in S06B-Bj and USDA 76-Be carried a *repABC* operon encoding all proteins required for autonomous replication (56). In fact, *repABC* plasmids are common in Alphaproteobacteria including the genera *Bradyrhizobium* and *Rhizobium* (56), suggesting these plasmids may have a broad host range. Conjugal transfer (*tra*) and type 3 secretion system (*T3SS*) operons on the SB plasmids suggest they are mobilized via conjugation (57). Both S06B-Bj and USDA 76-Be plasmids carried 58% and 45% of the total number of IS observed in their respective genomes (Fig. 4) increasing the chances of gene transfer between the bacterial chromosome and the plasmid (58). These plasmids also carried several metabolic genes including nodulation factors integrated between IS elements. For example, two oxygen-sensitive class III ribonucleotide triphosphate reductase (RNR) genes (59) were found between IS in pS06B2 (location:2,744–9,280 and 261,129– 267,596 bp). Interestingly, only oxygen-dependent RNRs were present in the SB chromosome and the class III RNR acquisition may be useful for SB in the oxygen limited conditions prevalent in soybean root nodules (60).

In addition to IS and plasmids, bacteriophages also contribute to the bacterial mobilome (51). Numerous studies have addressed the roles of IS and plasmids as mobile elements in SB, but to the best of our knowledge spontaneously produced phage (SPP) have not been previously reported. Our discovery of SPP adds to the existing repertoire of mobile elements in SB. Coverage analysis of phage reads shows that these regions are involved in specialized and generalized transduction events with a potential to carry genes over large distances not possible via IS or plasmids. The IS elements present in USDA 122-Bd prophage further intertwine prophages and the rest of the mobilome (Supplementary Fig. S3).

TEM analysis showed that all the SPP were tailed phage, and large terminase protein (TerL) analysis suggested they have different genome packaging mechanisms (Fig. 8). TerL proteins identify specific pac (headful packaging) and cos (cos-type packaging) sites on the phage genomes (38). Pac- and cos-like sites that randomly occur in bacterial genomes are sometimes packaged by phage terminases. Additionally some have argued that headful packaging phages have a propensity for generalized transduction (61). While further studies are needed to confirm specialized transduction, mapping analysis of SPP ppUSDA76Be-2 and ppS10JBj-1 suggested that headful packaging terminases could be involved in specialized transduction in these SPP.

Bradyrhizobia IS and plasmids have been shown to transfer symbiotic genes between different strains of bradyrhizobia (27, 28). While such transfers can have a significant impact on their gene pool, they require direct contact between two bradyrhizobia cells. The SPP identified and sequenced in this study are a strong indication that bradyrhizobia could use phages as a mechanism for HGT. Unlike IS, plasmids, and direct DNA transformation events, phages do not require cell to cell contact and are relatively stable in environmental conditions, thereby potentially increasing gene transfer events.

## Materials and Methods

### Bradyrhizobium culture and growth conditions

S06B-Bj, S10J-Bj, USDA 122-Bd, and USDA 76-Be strains were stored in 25% glycerol at -80°C and grown on modified arabinose gluconate (MAG) agar (ATCC medium 2233) to obtain single colonies. Bacterial cultures were grown from these single colonies in MAG broth 3–5 d at 30 °C with shaking at 200 rpm.

### Temporal dynamics of spontaneous phage production

SB cultures were inoculated from 3–5 d old starter cultures and grown in 50 mL MAG broth. At 0, 6, 12, 24, 36, and 48 h post-inoculation, 1 mL and 0.5 mL samples were collected for bacteria and virus enumeration, respectively. Cells in the 1 mL samples were counted using a hemocytometer. Cells were removed from the 0.5 mL samples by centrifugation at 5,000 × g for 5 min and subsequent filtration of the supernatant with a 0.22 µm Whatman Anotop syringe filter (Millipore Sigma, Burlington, MA). The filtrate was fixed with a final concentration of 0.1% formalin. Fixed phage-like particles were captured on a 0.02 µm Whatman Anodisc filter (Millipore Sigma), stained with 2.5X SYBR® Gold (Thermo Fisher Scientific, Waltham, MA), and counted using epifluorescence microscopy (62).

### Transmission electron microscopy of spontaneously produced phages

Spontaneously produced phages were isolated for transmission electron microscopy (TEM) from turbid SB cultures. Cells were removed by centrifugation at 5,000 × g for 5 min followed by supernatant filtration through a 0.22 µm Whatman Anotop syringe filter. Viruses in the filtrate were concentrated using 100 kDa Amicon filters (Millipore Sigma) and stored in MSM buffer at 4 °C (63). Phage particles were negatively stained with 2% uranyl acetate and TEM imaged (Zeiss Libra 120).

### Isolation and sequencing of bacterial DNA

DNA was isolated from stationary phase SB cultures using the AllPrep PowerViral DNA/RNA kit (Qiagen, Germantown, MD) following the manufacturer’s protocol. Twenty kilobase pair SMRT-bell sequencing libraries were constructed from 5–10 µg of DNA and sequenced using the PacBio RS II sequencer.

Long-read PacBio RS II sequence data from S06B-Bj, USDA 122-Bd, and USDA 76-Be were assembled using a hierarchical genome assembly process with HGAP2 SMRT Analysis Server v2.3.0. S10J-Bj was assembled using HGAP4 via SMRT Link v9.0 with the Falcon override option (cfg overrides pa_dbsplit_option = - x500 -s200). All genomes were polished using Quiver. Circlator (64) and BLAST (65) analyses were performed to check for genome circularization. CheckM v 1.15 (66) was used to assess genome completion and contamination.

### Bacterial genome and mobilome annotation

Assembled bacterial genomes were annotated using Rapid Annotation using Subsystem Technology (RAST v2) (67) and Prokka v1.14 (68). Presence and organization of nodulation (31), nitrogen fixation (31), and rhizobitoxine islands (69) was manually curated.

Multiple bioinformatic approaches were used for predicting the mobilome (insertion sequences, plasmids, and prophages) of each genome. Insertion sequence elements were identified using ISEscan (70). Candidate plasmids were identified through screening for a *repABC* operon for origin of replication, *tra* operon for conjugation, and *par* operon for partition genes. Putative plasmid contigs were confirmed using the PLSDB (37) database (max p value: 0.1, min identity: 0.9, winner takes all strategy). PhiSpy v4.3 (71), PHASTER (72), and Prophinder v0.4 (73) were used for predicting and annotating prophage regions.

### Isolation, sequencing, assembly, and mapping of phage DNA

Phages produced in USDA 76-Be cultures were isolated and concentrated using a modified iron chloride flocculation method (74). Cells were pelleted from 1 L of a 3.5 d old USDA 76-Be culture with centrifugation at 10,000 × g for 10 min, washed with fresh MAG media, and used for bacterial DNA extraction. The supernatant was filtered through 0.22 µm Stericup filters (Millipore Sigma), removing remaining bacterial cells, and pH adjusted to 7.5. Phages within the 0.22 µm filtered supernatant were flocculated by adding 1 mL of a 10 g/L FeCl_3_ stock solution per liter of media, pelleted by centrifugation at 4,000 × g for 15 min, and resuspended in 10 mL of Ascorbate-EDTA buffer. The phage suspension was centrifuged at 4,000 × g for 10 min and washed twice with 5 mL MSM buffer in Centricon Plus-70 100 kDa centrifugal filters (Millipore Sigma). Phage concentrates were stored in 200 µL MSM buffer at 4 °C. Iron chloride method yielded low amount of phage DNA.

Five hundred milliliters of a 3.5 day old bradyrhizobia culture was serially filtered through Pellicon XL 50 Biomax 0.22 µm and 100 kDa ultrafiltration modules (Millipore). The 0.22 µm retentate was recirculated three times to increase virus flowthrough to the 100 kDa filter. Cells were pelleted from 0.22 µm retentate by centrifugation at 10,000 × g for 10 min, washed with fresh MAG media, and used for bacterial DNA isolation. The 100 kDa retentate was spun through Centricon Plus-70 centrifugal filter, and the resulting phage concentrates were washed with 5 mL MSM buffer using Amicon 100 kDa filters. Any remaining bacterial DNA contamination in phage concentrates was removed using the DNase-I kit (Ambion, Austin, TX) with 30 °C incubation. A 16S PCR assay was used for confirming the absence of bacterial DNA contamination (75). Phage concentrates were stored in MSM buffer at 4 °C.

DNA from the cellular pellets (*i.e*., the host genome control) and phage concentrates was isolated using the AllPrep PowerViral DNA/RNA kit (Qiagen) following the manufacturer’s recommended protocol. DNA libraries were constructed using the Nextera DNAFlex library kit (Illumina, San Diego, CA) according to the manufacturer’s instructions. Libraries were sequenced using Illumina MiSeq (2 * 101 bp; USDA 76-Be) or Illumina NextSeq 550 (2 * 101 bp; S06B-Bj, S10J-Bj, and USDA 122-Bd). An in-house wrapper script (https://github.com/mooreryan/qc) based on Trimmomatic (76) and FLASH (77) was used to process the Illumina reads. Bowtie2 (78) was used to map host genome control and phage reads to their respective host HGAP-assembled genomes. Phage reads were assembled using CLC genomics workbench v20.0 using default settings.

### Phylogenetic analysis of terminase (TerL) large subunit sequences

Replication strategies and DNA packaging mechanisms can be predicted by phylogenetic analysis of the phage terminase large subunit (TerL) peptide sequence. TerL proteins from SB phages, S06B-Bj, S10J-Bj, USDA 76-Be, USDA 122-Bd were aligned with 242 phage TerL protein sequences (Supplementary data S2) from the UniProtKb/SwissProt and NCBI Virus databases using MAFFT (79) (FTT-NS-i X2 algorithm, default settings). An unrooted phylogenetic tree was constructed with FastTree (80) (default setting) in Geneious v10.2.3 (81). Conserved domains were determined by using RPS-BLAST (82) against the Pfam v32 (83) database and mapped onto the phylogenetic tree using Iroki (84).

## Acknowledgements

This research work was funded by the National Science Foundation under Grant No. 1736030. We thank University of Delaware Bioinformatics Core Facility for computational hardware, software used for analysis, and BIOMIX computational cluster. We thank the University of Delaware Sequencing and Genotyping Centre (USDGC) core for assistance with sequencing.We thanks University of Delaware Bioimaging Center for assistance with TEM imagining. Use of these facilities and microscopy access was supported by grants from the NIH-NIGMS (P20 GM103446), the NIGMS (P20 GM139760), the State of Delaware, and Delaware Biotechnology Institute.

## Data availability

All the data generated in this paper are part of an umbrella Bioproject- PRJNA686080. Specifically, genome and SRA for individual isolates are available in the following Bioprojects: S06B-Bj - PRJNA686124, S10J-Bj - PRJNA686125, USDA 122- Bd - PRJNA686127, and USDA-76-Be - PRJNA686128.

