## Supplementary Fig. S1 for "Spontaneously produced lysogenic phages are an important component of the soybean *Bradyrhizobium* mobilome"

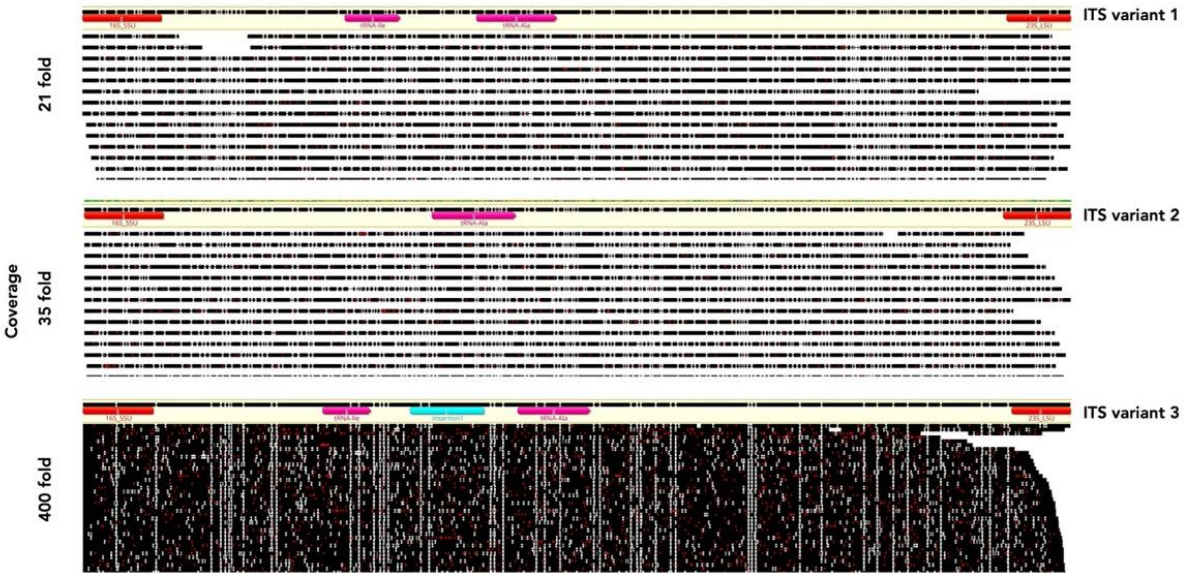


Supplementary Figure S1: Multiple internal transcribed spacer (ITS) variants were observed in the S10J-Bj genome. PacBio reads were mapped against the three ITS variants previously observed in S10J-Bj genome. While the final S10J-Bj assembly suggested the presence of two identical copies of ITS variant 3, PacBio read mapping suggested all three variants were present. However, read mapping coverage of ITS variant 1 (21 X) and ITS variant 2 (35 X) were low compared to ITS variant 3 (400 X).
