## Supplementary Fig. S2 for "Spontaneously produced lysogenic phages are an important component of the soybean *Bradyrhizobium* mobilome"

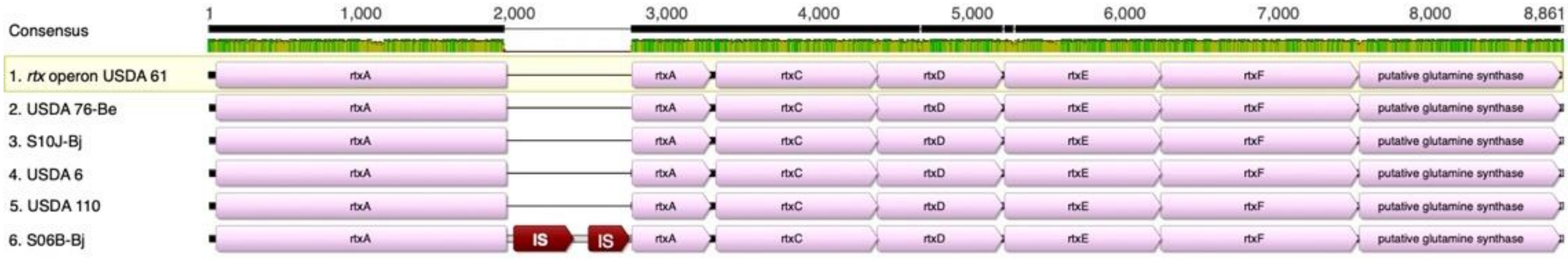


Supplementary Figure S2: Truncated *rtx* operons observed in non-rhizobitoxine producing bradyrhizobia genomes. A 9,000 bp region with >80% similarity (100% length) to the reference strain USDA 61 rhizobitoxine operon was found in USDA76-Be, S06B-Bj, and S10J-Bj genomes. Similar regions were also observed in reference strains USDA 6 and USDA 110. Both *B. elkanii* strains, USDA 76-Be and rhizobitoxine-producing USDA 61, had intact *rtx* operons, while the operon was truncated by premature stop codons in other strains. Additionally, IS were integrated on the *rtxA* gene in S06B-Bj.
