## Supplementary Fig. S3 for "Spontaneously produced lysogenic phages are an important component of the soybean *Bradyrhizobium* mobilome"

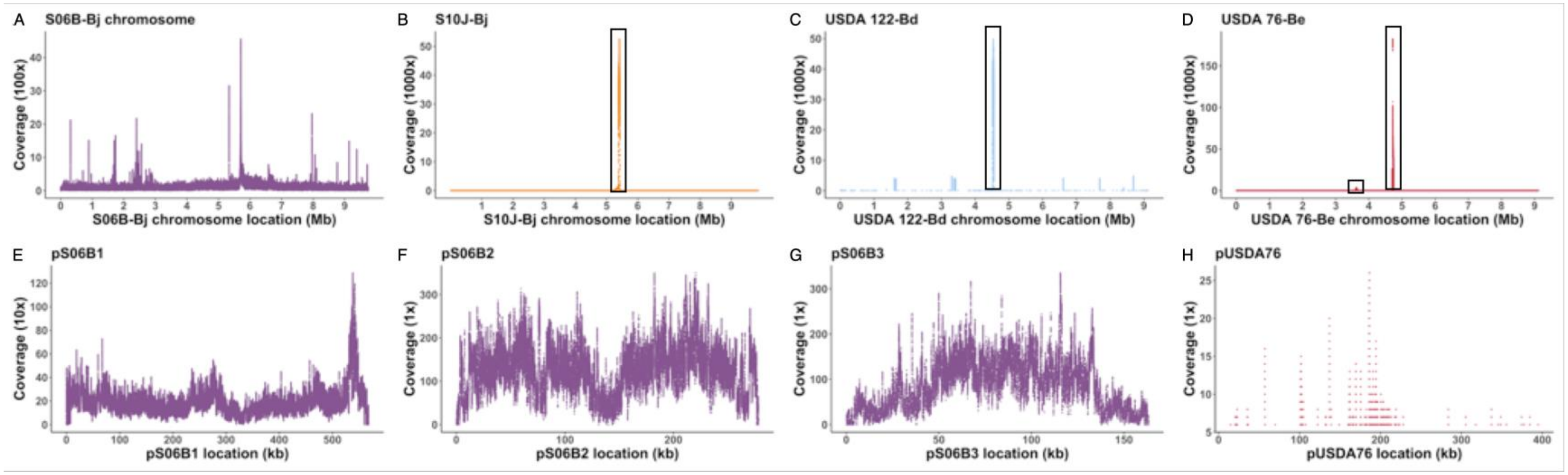


Supplementary Figure S3: Mapping of phage reads to bacterial chromosomes and plasmids. Sequence reads from spontaneously produced phages mapped to bradyrhizobia chromosomes (A-D) and plasmids (E-H). Phages reads showed enriched mapping (boxed) for B) S10J-Bj, C) USDA 122-Bd, and D) USDA 76-Be. There were no phage specific enriched regions observed for S06B-Bj or any of the plasmids. USDA 122-Be and S06B-Bj had several regions of high coverage which corresponded to insertion sequences. Prophage ppUSDA122Bd-1 contained an insertion sequence, homologs of which were present at multiple locations on the USDA 122-Bd bacterial chromosome. Mapping of insertion sequence reads resulted in additional high coverage regions outside of the prophage region. IS read mapping outside of the prophage region resulted in decreased coverage for that insertion sequence in ppUSDA122Bd-1 (Fig. 7).
