## Supplementary Fig. S4 for "Spontaneously produced lysogenic phages are an important component of the soybean *Bradyrhizobium* mobilome"

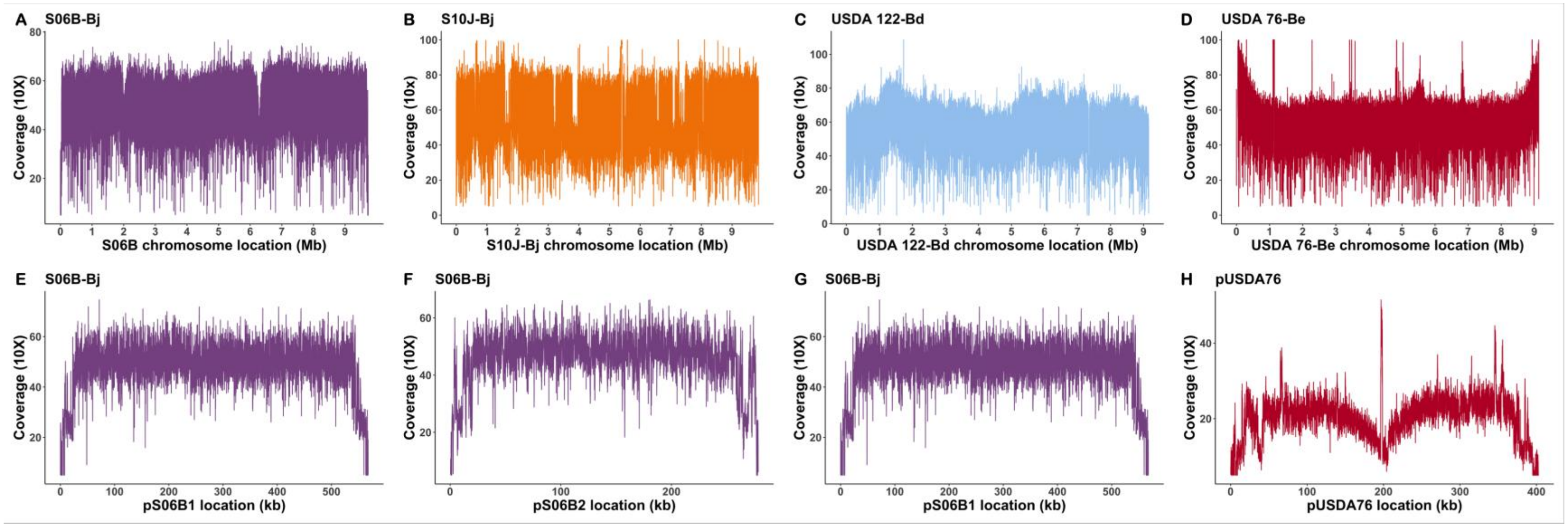


Supplementary Figure S4: Mapping of bacterial host DNA sequencing reads to their respective chromosome. Sequence mapping plots against chromosomes: A) S06B-Bj; B) S10J-Bj; C) USDA 122-Bd; and D) USDA 76-Be, and plasmids: E) pS06B1; F) pS06B2; G) pS06B3; and H) pS06B4. Each of these cultures spontaneously produced prophages. Mapping resulted in even coverage ranging from 100–1000x for chromosomes and 400-600x for plasmids. Read mapping of phage DNA isolated from these cultures showed coverage values 10–100 fold higher than those observed for the bacterial DNA sequences (Fig. 7, Fig. S3).
